# Evaluation of a novel isothermal microcalorimetry-based sterility test

**DOI:** 10.64898/2026.02.11.705057

**Authors:** Indra Sioen, Tom Coenye

## Abstract

Parenteral drugs must meet strict release criteria to ensure patient safety upon administration. Sterility is a critical requirement, and is typically assessed using the compendial U.S. Pharmacopoeia (USP) <71> sterility test. However, with the growing demand to reduce batch release times, the test’s 14-day incubation period is becoming a concern, emphasizing the need for more rapid alternatives. Here, we compared the performance of the calScreener+ isothermal microcalorimetry (IMC) device (Symcel) to that of the compendial USP <71> sterility test, using a panel of sixteen microorganisms (six USP <71> reference strains and ten field isolates) in two inoculum sizes (100 and 5 CFU). The IMC-based method detected a higher number of positive samples compared to the compendial method (95.8% vs 87.5%; p < 0.05). Furthermore, IMC was consistently faster, reducing mean detection times from 43 hours to 19 hours at 100 CFU and from 46 hours to 28 hours at 5 CFU (p < 0.001). In conclusion, the calScreener+ IMC device shows promise as a rapid and sensitive alternative to the compendial sterility test, with the potential to speed up batch releases without compromising patient safety.

## Introduction

Sterility is a critical quality attribute for parenteral drugs, and regulatory authorities such as the U.S. Food and Drug Administration therefore mandate sterility testing prior to batch release **(1)**. The current gold standard, described in the U.S. Pharmacopoeia (USP) <71>, is a presence/absence test in which samples are inoculated into two culture media (Fluid Thioglycolate Medium [FTM] and Tryptone Soy Broth [TSB]) using either membrane filtration or direct inoculation, and then incubated for at least 14 days with periodic visual inspection for turbidity **(2)**. The test was originally intended for bulk-manufactured sterile pharmaceuticals, and has been used essentially unchanged since its introduction in 1932 **(3)**. However, the long incubation period and reliance on (subjective) visual readouts make it poorly suited for emerging biological therapies, that often have a short shelf-life and/or are intrinsically turbid, and where rapid administration can be crucial for successful therapy **(4, 5)**. Consequently, these biologics are frequently infused in the patient before the final sterility assessment, or are frozen until completion of the 14-day protocol, introducing logistical challenges and impairing manufacturing efficiency **(6, 7)**. Therefore, there is a growing demand for rapid microbiological methods that can be applied for both end-product release testing and in-process controls **(8-11)**. Regulatory authorities have also issued guidelines to support their validation **(12, 13)**.

Several automated rapid technologies have been evaluated for sterility assessment. Established growth-based methods include respiration-based systems (e.g. BACT/ALERT 3D, BD BACTEC) and adenosine triphosphate (ATP) bioluminescence measurement systems (e.g. Milliflex Rapid System); use of such systems can reduce the time to detection (TTD) to 4-7 days **(9, 14-18)**. While this is a significant improvement, these incubation times are still long for drugs with a short shelf-life. For these drugs, an 48-hour, overnight or even real-time microbial test would be more appropriate to support real-time product release **(13)**. Additional limitations include reported difficulties of some rapid alternatives in detecting slow-growing mold species and bacteria, and/or the destructive nature of some tests, which precludes downstream analysis (e.g. microorganism identification) **(10, 11)**. This emphasizes the need for other approaches based on alternative detection principles (such as other growth-based parameters).

Heat is a byproduct of the metabolism of microorganisms, and isothermal microcalorimetry (IMC) devices allow to continuously measure this metabolic heat production in real-time. As IMC measures a fundamental physiological parameter, the method is inherently label-free and suitable for diverse sample types, including turbid, viscous or solid samples. In addition, IMC is a non-destructive method, allowing downstream analysis of the sample. Moreover, several studies have shown that IMC can detect microorganisms earlier than other growth-based methods while maintaining a high sensitivity **(19, 20)**. In the present study, we evaluated the performance of the calScreener+ IMC device (Symcel) and compared it against the USP <71> compendial sterility test, by comparing the number of positive cultures detected and the TTD.

## Materials and methods

### Microbial strains and culture conditions

Sixteen microorganisms (nine Gram-positive bacteria, three Gram-negative bacteria, three molds and one yeast) were used in this study (Table 1). These include organisms recommended for growth promotion testing of the media included in USP <71> and isolates recovered from pharmaceutical products and pharmaceutical production environments **(2)**. Pure cultures and overnight cultures were maintained on culture media and under culture conditions described in Table 1.

**Table 1.** List of microorganisms used, with culture conditions.

| Species | Strain designation | Additional information | Media and culture conditions |
| --- | --- | --- | --- |
| *Acidovorax delafieldii* | LPM-S1 | Isolated from pharmaceutical production environment (Belgium) | Tryptone Soy Agar (TSA)<sup>a</sup>; Nutrient Broth (NB)<sup>a</sup><br>Aerobic<sup>b</sup> |
| *Aspergillus brasiliensis* | ATCC 16404 | USP <71> reference strain | Sabouraud Dextrose Agar (SDA)<sup>a</sup><br>Aerobic<sup>b</sup> |
| *Aspergillus versicolor* | LPM-S2 | Isolated from pharmaceutical production environment (Belgium) | SDA<br>Aerobic<sup>b</sup> |
| *Bacillus spizizenii* | ATCC 6633 | USP <71> reference strain | TSA; NB<br>Aerobic<sup>b</sup> |
| *Bacillus subtilis* | LPM-S3 | Isolated from pharmaceutical production environment (Belgium) | TSA; NB<br>Aerobic<sup>b</sup> |
| *Burkholderia contaminans* | LPM-S4 | Isolated from contaminated liquid docusate | TSA; NB<br>Aerobic<sup>b</sup> |
| *Candida albicans* | ATCC 10231 | USP <71> reference strain | SDA, Sabouraud Dextrose Broth<sup>c</sup><br>Aerobic<sup>b</sup> |
| *Clostridium sporogenes* | ATCC 19404 | USP <71> reference strain | Columbia Agar supplemented with 5% sheep blood (CBA)<sup>a</sup>; Reinforced Clostridial Medium (RCM)<sup>c</sup><br>Anaerobic<sup>b</sup> |
| *Cutibacterium acnes* | LMG 16711 | Isolated from human skin (patient with facial acne) | CBA; RCM Anaerobic<sup>d</sup> |
| *Micrococcus luteus* | LPM-S5 | Isolated from human saliva | CBA; NB Aerobic<sup>b</sup> |
| *Penicillium chrysogenum* | LPM-S6 | Isolated from pharmaceutical production environment (Belgium) | SDA Aerobic<sup>b</sup> |
| *Pseudomonas paraeruginosa* | ATCC 9027 | USP <71> reference strain | TSA; NB Aerobic<sup>b</sup> |
| *Staphylococcus aureus* | ATCC 6538 | USP <71> reference strain | TSA; NB Aerobic<sup>b</sup> |
| *Staphylococcus epidermidis* | LPM-S7 | Isolated from pharmaceutical production environment (Belgium) | TSA; NB Aerobic<sup>b</sup> |
| *Staphylococcus hominis* | LPM-S8 | Isolated from pharmaceutical production environment (Belgium) | TSA; NB Aerobic<sup>b</sup> |
| *Streptococcus pyogenes* | LMG 14237 | Isolated from throat of a patient suffering from rheumatic fever | TSA; NB Aerobic<sup>b</sup> |

<sup>a</sup> Supplier: Neogen
<sup>b</sup> Overnight culture: 37°C, 16 hours
<sup>c</sup> Supplier: Lab M
<sup>d</sup> Overnight culture: 37°C, 40 hours

### Preparation of microbial suspensions

Overnight cultures were diluted in physiological saline (PS, 0.9% NaCl) to final theoretical concentrations of 100 and 5 CFU per 50 µL. For molds, spore suspensions were prepared directly from the Sabouraud Dextrose Agar (SDA, Neogen) cultures. 5 mL of PS supplemented with 0.1% Tween 20 (Sigma-Aldrich) was added to each plate to promote spore detachment. The resulting suspensions were collected, and spores were counted using a Neubauer-improved chamber (Marienfeld) and a Olympus B201 optical microscope (Olympus). The spore suspensions were subsequently diluted to final theoretical concentrations of 100 spores and 5 spores per 50 µL. To confirm the microbial load, 50 µL of each suspension was plated in duplicate via the pour plate method, using the same media and conditions described above. Following incubation, colonies were counted and the actual number of CFU was calculated. To reduce variability, both the compendial and IMC-based sterility test were inoculated in parallel from a single microbial suspension.

### Compendial sterility test

The compendial sterility test was performed using the direct inoculation method. Test tubes containing 5 mL of TSB (Neogen) (for *Acidovorax delafieldii, Aspergillus brasiliensis, Aspergillus versicolor, Bacillus spizizenii, Bacillus subtilis, Burkholderia contaminans, Candida albicans, Micrococcus luteus* and *Penicillium chrysogenum*) or FTM (Oxoid) (for *Clostridium sporogenes, Cutibacterium acnes, Pseudomonas paraeruginosa, Staphylococcus aureus, Staphylococcus epidermidis, Staphylococcus hominis* and *Streptococcus pyogenes*) were inoculated with 50 µL suspension and subsequently incubated at the temperatures recommended by the USP (25.0°C for TSB, 35.0°C for FTM; actual ranges from 24.7-25.1°C and 34.0-35.1°C, respectively) **(2)**. Although unlikely to have influenced the outcome, it should be noted that these temperatures fall outside the USP <71> recommended ranges by 0.1°C. Cultures were checked daily for 14 days for signs of turbidity. All experiments were carried out in biological triplicates.

### Microcalorimetry-based sterility testing with the calScreener+ device

For the calScreener+ sterility test, 350 µL of the appropriate growth medium (as described above) was injected into sterile glass vials (Symcel) using a 1 mL syringe (Becton Dickinson) with a 26G needle (Henke Sass Wolf), followed by the addition of 50 µL of the microbial suspensions. Thermodynamic reference vials were filled with 400 µL of PS. Vials were sealed and incubated in the calScreener+ instrument at 24.80°C ± 0.01°C (TSB) or 34.80°C ± 0.01°C (FTM) (these temperatures provide maximal baseline stability). Metabolic heat production was monitored in real time, generating thermograms that plot heat flow (µW) over time (calView2 software, Symcel). Samples were considered positive if the heat flow exceeded 1 µW. The timepoint at which this occurred, was recorded as the TTD. Suitability of the detection threshold was verified by assessing baseline characteristics of uninoculated medium samples (400 µL, in triplicates). Short-term noise (standard deviation [SD] of the heat flow signal over a representative one-hour interval) and signal stability (slope of the signal over 24 hours) were calculated to study both random fluctuations and long-term baseline stability, respectively. All IMC experiments were performed in biological triplicates (with each of these biological replicates consisting of a technical duplicate). A biological replicate was considered positive if one of two technical replicates exceeded the 1 µW detection threshold.

### Statistical analysis

The exact McNemar test was used to compare the number of positive cultures detected by the compendial and calScreener+ method for each inoculum concentration. TTD per inoculum size was compared between methods using a Wilcoxon signed rank test. To compare the TTD between methods across organisms, a paired sample t-test was performed after confirming normal distribution of the data (QQ plots). For the 100 CFU inoculum size, TTD comparisons per organism were performed with paired sample t-tests. Bonferroni correction was used to adjust for multiple comparisons. All statistical tests were performed using IBM SPSS Statistics (version 29.0.1.0). For all tests, a p value of < 0.05 was considered statistically significant. Replicates not detected within the 336-hour detection window result in right-censored data that cannot be analyzed by a standard parametric test (e.g. t-test). Although survival analysis methods (e.g. Cox regression) can handle censored data, the sample size per condition (n = 3) in this study does not provide sufficient statistical power. Therefore, statistical analysis was not performed for the 5 CFU inoculum size, as some replicates remained undetected with this inoculum.

## Results

### Confirmation of the challenge inoculum sizes

For the comparison of the two methods, challenge inocula theoretically containing 100 and 5 CFU were prepared. Plating showed that the actual values ranged from 250 to 22 CFU (mean: 62 CFU; SD: 47 CFU) and 19 to 1 CFU (mean: 4 CFU; SD: 3 CFU), respectively. Actual inoculum sizes are provided in Supplementary Table S1.

### Confirmation of microcalorimetric baseline stability

Thermograms from uninoculated TSB and FTM samples were analyzed to evaluate the baseline characteristics of the microcalorimetric system (Supplementary figure S1). Both culture media provided stable baseline values, with acceptable short term noise (0.019 µW in TSB and 0.041 µW in FTM) and baseline drift (0.00069 µW/h in TSB and -0.014 µW/h in FTM). Furthermore, mean and SD of the 24-hour baseline signal were calculated for both media. To distinguish true signal from baseline fluctuations, a commonly applied criterion is that the detection threshold should exceed the mean baseline value plus three times the SD. For TSB, mean signal + 3SD equals 0.162 µW; the 1 µW detection threshold approximately equals mean + 14SD. For FTM, mean + 3SD equals 0.164 µW; 1 µW approximately equals mean + 11SD. Combined, these data support the suitability of the 1 µW detection threshold to distinguish between baseline fluctuations and metabolic activity.

### Microcalorimetry detects higher number of positive cultures than the compendial method

When using the higher inoculum size (theoretically 100 CFU), both methods detected the presence of all organisms tested. At the lower inoculum size (theoretically 5 CFU), the microcalorimetry-based method yielded more positive results than the compendial method (91.7% compared to 75.0%; p < 0.05) (Table 2). When considering both inoculum sizes together, the microcalorimetry-based method also detected more samples than the compendial test (95.8% vs. 87.5%; p < 0.05) (Table 2). However, as the true contamination state of the samples was not confirmed, these results should not be interpreted as evidence of superior sensitivity. Representative examples of thermograms obtained with different inoculum sizes for different organisms are shown in Figure 1. Additional thermograms are available upon request.

**Table 2.** Number of positive cultures detected by the compendial sterility test and the microcalorimetry-based method for the different inoculum sizes.

| Target inoculum size | Number of positive cultures / total number of cultures tested (%) |  |
| --- | --- | --- |
|  | Compendial method | Microcalorimetry-based method |
| 100 CFU | 48/48 (100%) | 48/48 (100%) |
| 5 CFU | 36/48 (75.0%) | 44/48 (91.7%)<sup>a</sup> |
| Total | 84/96 (87.5%) | 92/96 (95.8%)<sup>a</sup> |

<sup>a</sup> Statistically significant difference compared to compendial method; $p < 0.05$ (Exact McNemar test)

**Figure 1.**
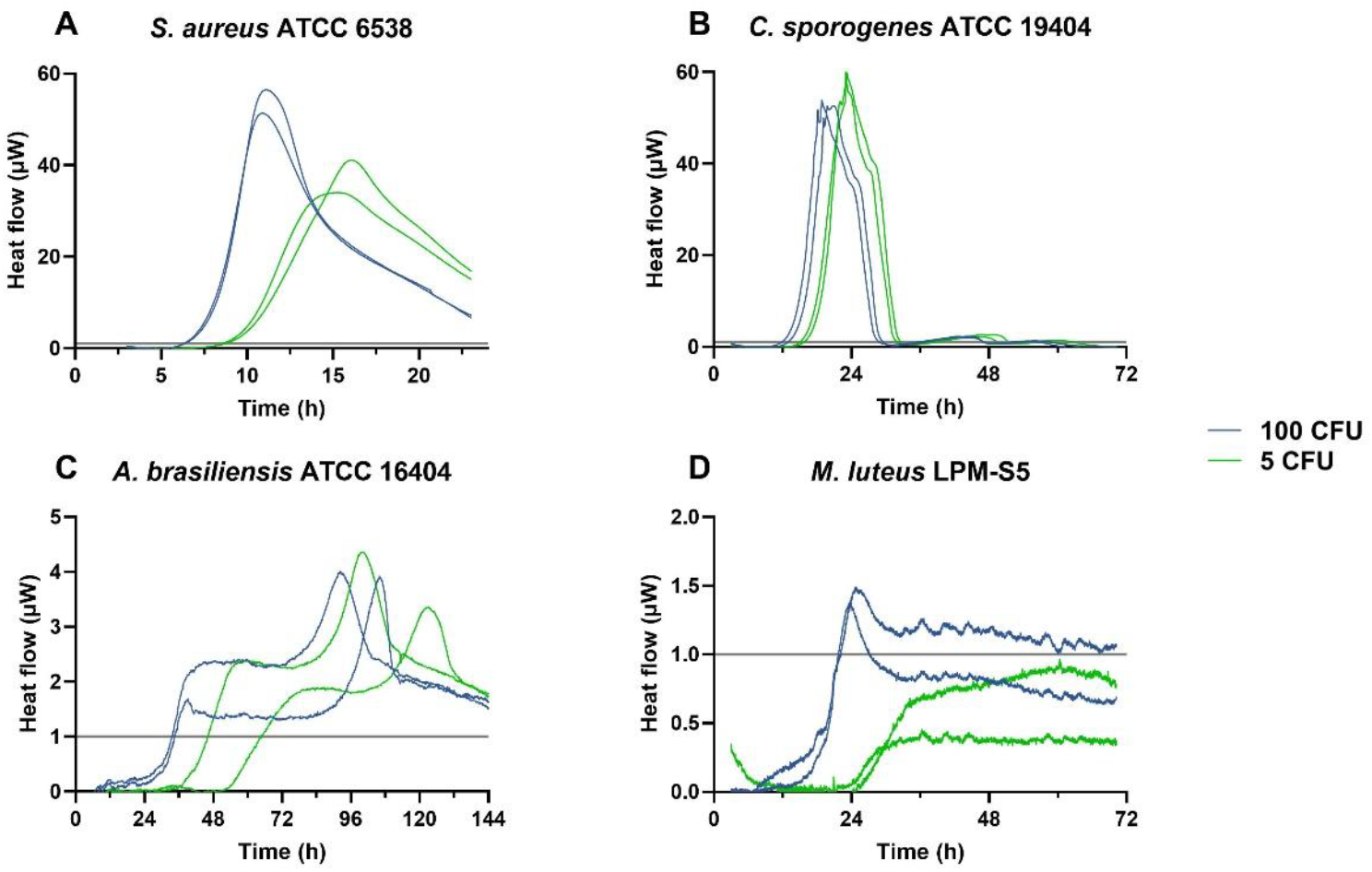
Thermograms for a biological replicate (consisting of two technical replicates) of (A) *S. aureus* (in FTM), (B) *C. sporogenes* (in FTM), (C) *A. brasiliensis* (in TSB) and (D) *M. luteus* (in TSB, lower inoculum size undetected) for the theoretical inoculum sizes of 100 CFU and 5 CFU. The grey line at 1 µW indicates the detection threshold.

### Microcalorimetry may shorten the time to detection

Detection times for each of the sixteen organisms were determined in biological triplicate using both the compendial and calScreener+ sterility tests. The mean TTD was then calculated for all samples and per organism.

At both inoculum sizes, the calScreener+ detected the presence of microorganisms more rapidly than the compendial method (Figure 2). Mean detection times were shortened by 24 hours for 100 CFU and by 18 hours for 5 CFU (Wilcoxon signed rank test, p < 0.001). Furthermore, all sixteen organisms were detected more rapidly, with statistically significant differences between average TTD for almost all organisms at 100 CFU (Table 3; Figure 3). The most pronounced difference in TTD (although not significant after correction for multiple testing) was observed for *C. albicans*, that was detected on average two days earlier using microcalorimetry. Similarly, the presence of the filamentous fungi *A. brasiliensis, A. versicolor* and *P. chrysogenum* and the Gram-negative bacterium *B. contaminans* was detected at least one day sooner using microcalorimetry than with the compendial method (Table 3; Figure 3).

**Table 3.**
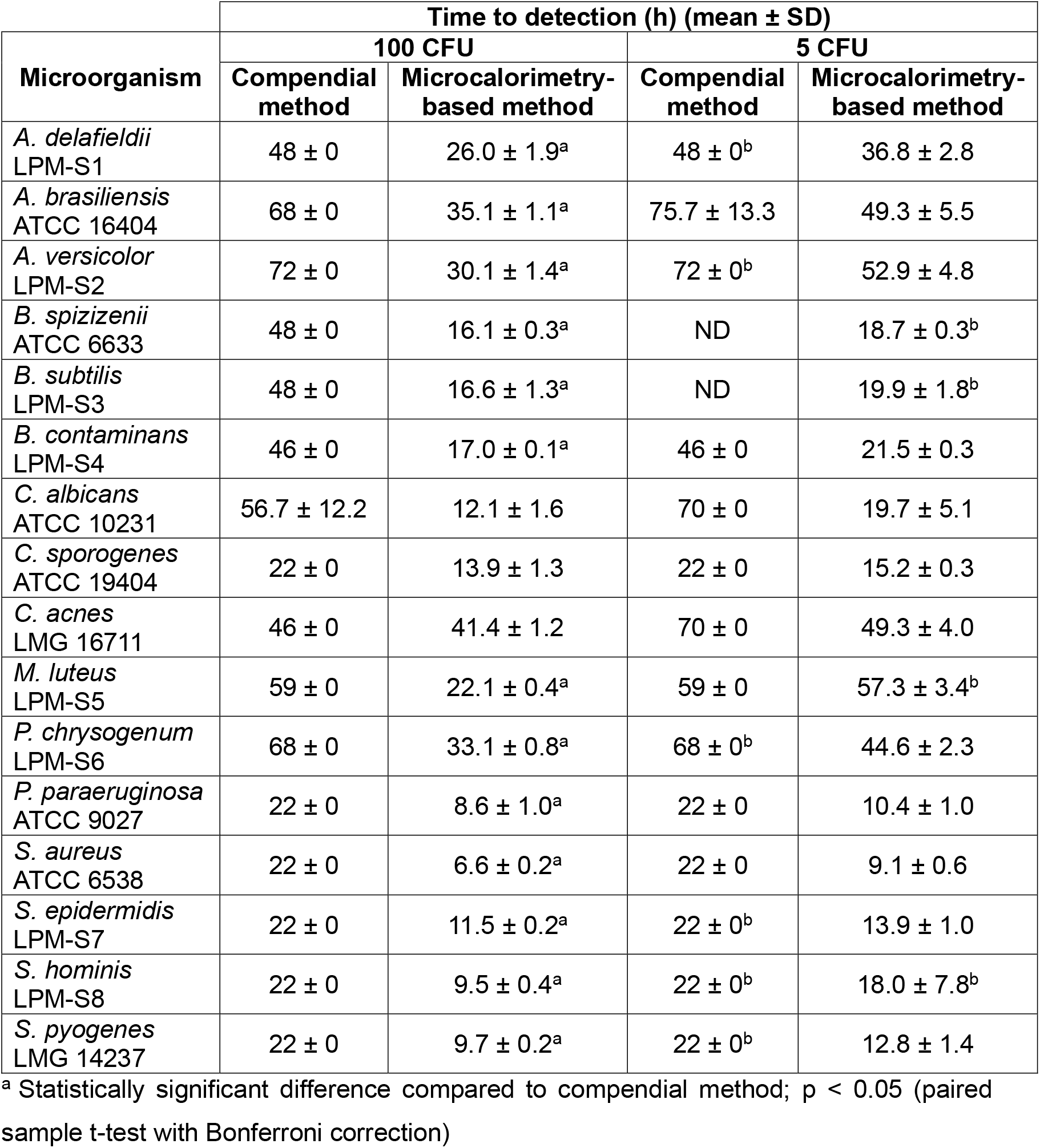
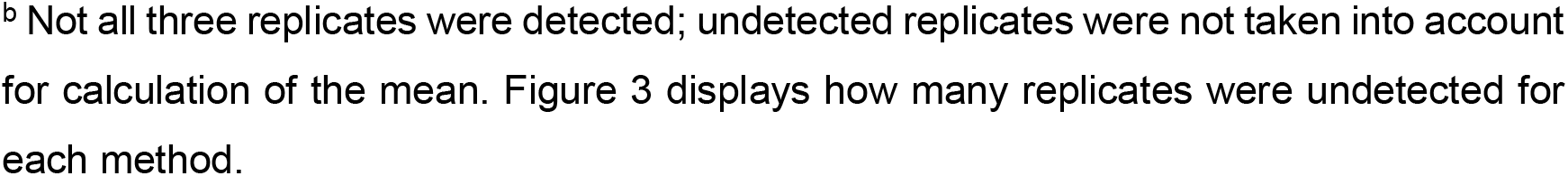
Detection times for each method for each organism. Data are presented as mean ± standard deviation (SD, n = 3). ND, not detected (i.e. all three replicates were negative). Statistics were performed on the 100 CFU inoculum size only (as for these, there are no missing data).

| Microorganism | Time to detection (h) (mean ± SD) | | | |
| --- | --- | --- | --- | --- |
|  | 100 CFU |  | 5 CFU |  |
|  | Compendial method | Microcalorimetry-based method | Compendial method | Microcalorimetry-based method |
| *A. delafieldii*<br>LPM-S1 | 48 ± 0 | 26.0 ± 1.9<sup>a</sup> | 48 ± 0<sup>b</sup> | 36.8 ± 2.8 |
| *A. brasiliensis*<br>ATCC 16404 | 68 ± 0 | 35.1 ± 1.1<sup>a</sup> | 75.7 ± 13.3 | 49.3 ± 5.5 |
| *A. versicolor*<br>LPM-S2 | 72 ± 0 | 30.1 ± 1.4<sup>a</sup> | 72 ± 0<sup>b</sup> | 52.9 ± 4.8 |
| *B. spizizenii*<br>ATCC 6633 | 48 ± 0 | 16.1 ± 0.3<sup>a</sup> | ND | 18.7 ± 0.3<sup>b</sup> |
| *B. subtilis*<br>LPM-S3 | 48 ± 0 | 16.6 ± 1.3<sup>a</sup> | ND | 19.9 ± 1.8<sup>b</sup> |
| *B. contaminans*<br>LPM-S4 | 46 ± 0 | 17.0 ± 0.1<sup>a</sup> | 46 ± 0 | 21.5 ± 0.3 |
| *C. albicans*<br>ATCC 10231 | 56.7 ± 12.2 | 12.1 ± 1.6 | 70 ± 0 | 19.7 ± 5.1 |
| *C. sporogenes*<br>ATCC 19404 | 22 ± 0 | 13.9 ± 1.3 | 22 ± 0 | 15.2 ± 0.3 |
| *C. acnes*<br>LMG 16711 | 46 ± 0 | 41.4 ± 1.2 | 70 ± 0 | 49.3 ± 4.0 |
| *M. luteus*<br>LPM-S5 | 59 ± 0 | 22.1 ± 0.4<sup>a</sup> | 59 ± 0 | 57.3 ± 3.4<sup>b</sup> |
| *P. chrysogenum*<br>LPM-S6 | 68 ± 0 | 33.1 ± 0.8<sup>a</sup> | 68 ± 0<sup>b</sup> | 44.6 ± 2.3 |
| *P. paraeruginosa*<br>ATCC 9027 | 22 ± 0 | 8.6 ± 1.0<sup>a</sup> | 22 ± 0 | 10.4 ± 1.0 |
| *S. aureus*<br>ATCC 6538 | 22 ± 0 | 6.6 ± 0.2<sup>a</sup> | 22 ± 0 | 9.1 ± 0.6 |
| *S. epidermidis*<br>LPM-S7 | 22 ± 0 | 11.5 ± 0.2<sup>a</sup> | 22 ± 0<sup>b</sup> | 13.9 ± 1.0 |
| *S. hominis*<br>LPM-S8 | 22 ± 0 | 9.5 ± 0.4<sup>a</sup> | 22 ± 0<sup>b</sup> | 18.0 ± 7.8<sup>b</sup> |
| *S. pyogenes*<br>LMG 14237 | 22 ± 0 | 9.7 ± 0.2<sup>a</sup> | 22 ± 0<sup>b</sup> | 12.8 ± 1.4 |

<sup>a</sup> Statistically significant difference compared to compendial method; p < 0.05 (paired sample t-test with Bonferroni correction)
<sup>b</sup> Not all three replicates were detected; undetected replicates were not taken into account for calculation of the mean. Figure 3 displays how many replicates were undetected for each method.

**Figure 2.**
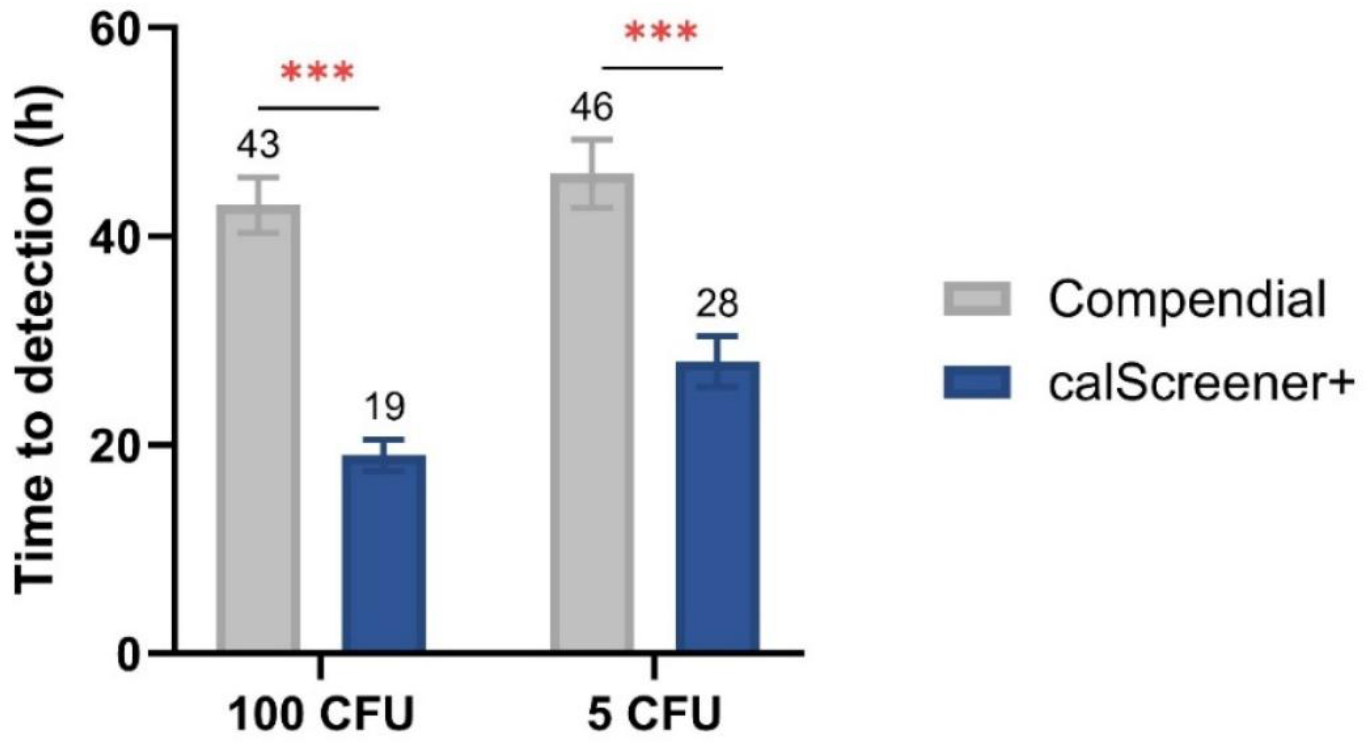
Mean detection times for each method, grouped per inoculum size. Error bars represent standard error of the mean. *** indicates p < 0.001 (Wilcoxon signed rank test).

**Figure 3.**
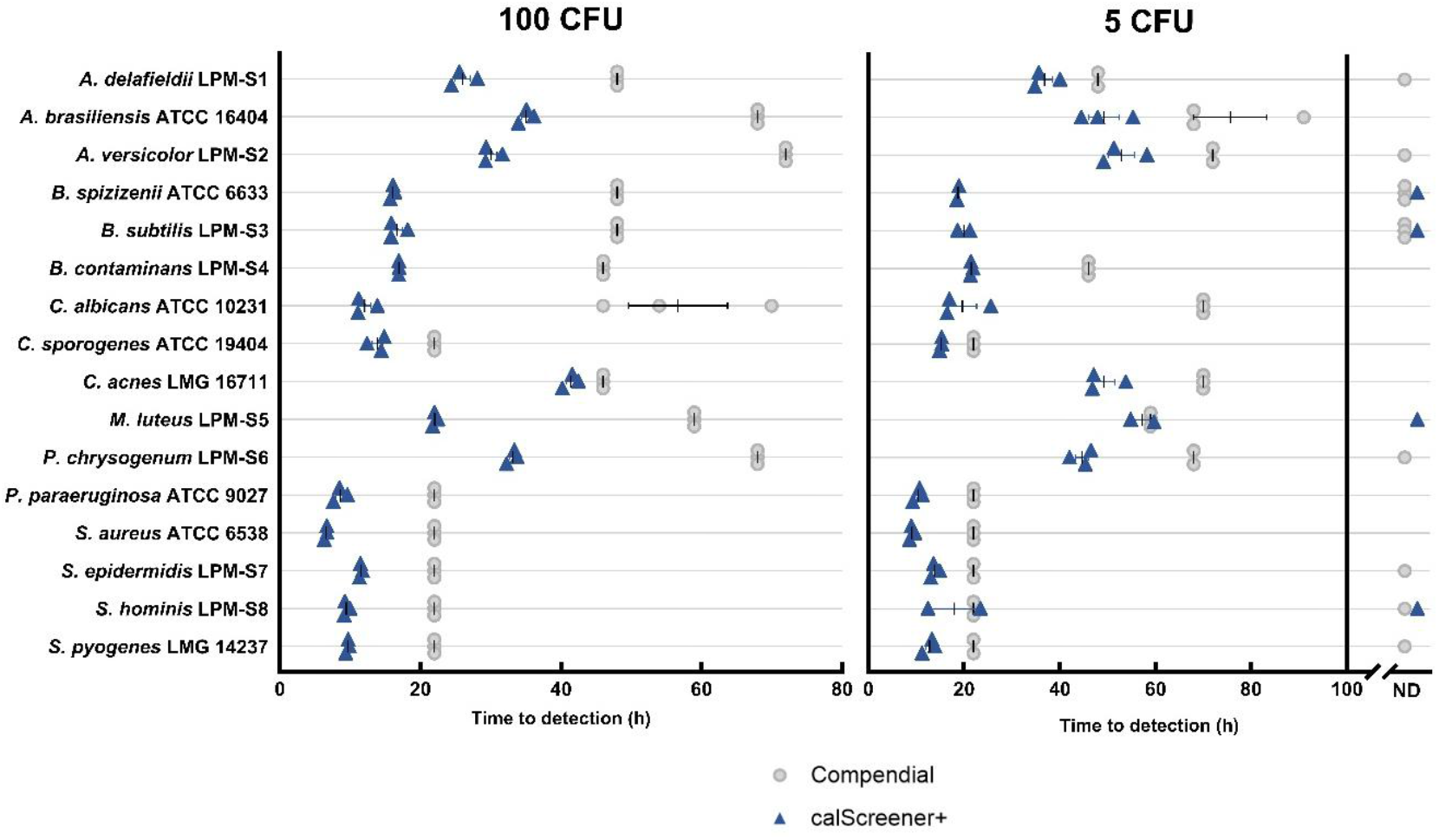
Time to detection for the individual replicates of the sixteen tested microorganisms for the 100 CFU inoculum (left) and the 5 CFU inoculum (right). Replicates for which no growth was observed within the 336 hour detection window are shown as not detected (ND). Error bars represent mean ± standard error of the mean.

### At the low inoculum size some replicates are not detected

TTD data for individual replicates are shown in Figure 3. At the low target inoculum size (5 CFU), 16.7% of all experiments (IMC method and compendial method combined) did not yield positive results within the 336-hour detection window. For both *Bacillus* spp. included, the calScreener+ yielded positive results in two out of three replicates, while no samples turned positive using the compendial method. For 12 other organisms, all replicates were detected by microcalorimetry, highlighting it’s potential to detect low-level contamination by various organisms (in this case *A. delafieldii, A. brasiliensis, A. versicolor, B. contaminans, C. albicans, C. sporogenes, C. acnes, P. chrysogenum, P. paraeruginosa, S. aureus, S. epidermidis* and *S. pyogenes*). One replicate of *M. luteus* and *S. hominis* remained negative using microcalorimetry; in case of *M. luteus*, a clear increase in metabolic heat flow was observed, but it did not surpass the predetermined 1 µW detection threshold (Figure 1).

## Discussion

Advances in calorimetric sensitivity and the introduction of multi-channel calorimetric devices have led to a growing interest in IMC for various applications, including rapid detection of microorganisms in clinical and industrial contexts **(21-25)**. Brueckner et al. already demonstrated IMC’s potential for sterility testing using artificially contaminated monoclonal antibody formulations, reporting shorter mean detection times than with visual inspection **(26)**.

In the present study, we explored the possibility of using the calScreener+ device for sterility testing purposes and compared it with the compendial USP <71> method. Our findings show that the IMC-based method detected a higher number of positive cultures, especially at the lower inoculum size (95.8% vs. 87.5% overall; 91.7% vs. 75.0% for 5 CFU), while also consistently reducing mean detection times for the sixteen organisms tested (on average from 43 to 19 h and from 46 to 28 h for 100 CFU and 5 CFU, respectively; p < 0.001). TTD differences can likely be attributed to the sensitivity of the underlying detection principle. While the human eye typically requires concentrations of ≥ 10^7^ cells/mL to detect turbidity changes, IMC devices can detect metabolic heat from as few as 10^4^ cells/mL **(27, 28)**.

A limitation of the present study is that, while IMC data were collected continuously in real-time, visual inspection was not continuous, leading to a potential overestimation of the TTD for the compendial test. However, the large and consistent TTD differences observed in our study do suggest actual analytical advantages. Moreover, periodic inspection reflects industrial practice. Continuous monitoring offers a clear advantage in Good Manufacturing Practice (GMP) environments as it enables objective, electronically recorded readouts. These can be automated to trigger real-time alerts, allowing faster corrective and preventive actions in case of contamination.

At the lower inoculum size (theoretical 5 CFU), IMC detected more positive cultures than the compendial method. The largest discrepancy between methods was seen for the *Bacillus* spp., where two out of three replicates were detected with IMC, while all replicates remained negative with the compendial method. Although the latter has a theoretical limit of detection (LOD) of 1-3 CFU, previous studies have reported difficulties at these lower inoculum levels **(11, 13)**. Plate count data for these strains suggested inoculum sizes around this theoretical LOD (2 CFU ± 1, Supplementary table S1). More extensive LOD studies could help to better understand the impact of low inoculum levels on both methods. Notably, variability in growth detection related to medium producer and batch have also been reported before, highlighting the importance of testing the growth-supporting properties of media; this was not performed in the present study **(25)**. IMC did fail to detect one replicate of the obligate aerobe *M. luteus*. Overall, this organism showed low signals, fluctuating around the 1 µW detection threshold. A potential explanation for this is that, in the current setup, the calScreener+ vials were fully filled with 400 µL of sample, leaving little headspace and thus possibly insufficient residual oxygen to support robust growth of this organism. Optimization could include a reduction of the total volume in order to increase oxygen availability in the vial.

16.7% of samples at the lower inoculum size did not become positive. When working with low inoculum sizes, the actual number of viable cells that are inoculated will vary and may deviate from the calculated inoculum. This variation arises from the random nature of cell distribution and can be estimated using a Poisson distribution. At 4 CFU (mean of the lower inoculum size), this means approx. 2% of samples will contain no viable cells and approx. 7% only a single cell, which might be insufficient to support detectable growth **(30, 31)**. Negative results observed in the present study are thus likely due to a combination of methodological limits and inherent stochastic effects. While these effects are inevitable, they can be reduced by increasing the inoculation volume and maintaining a tight control over the concentration of microorganisms in the suspension used for inoculation (e.g. by using Bioballs, bioMérieux). Furthermore, including a broader range of inoculum sizes (e.g. by testing a dilution series) could help to distinguish between results that are negative due to methodological limitations and those that are more likely to reflect stochastic under-inoculation.

In terms of experimental setup, every biological replicate in the microcalorimetric method consisted of two technical replicates, in line with standard practice and manufacturer recommendations. In contrast, the compendial method was performed using a single technical replicate. Technical duplicates are typically not included in compendial sterility testing. While our setup thus reflects routine practice for each method, it could partially explain the detection differences observed. Furthermore, our approach was designed to have the same number of CFU for both methods. As different volumes are used in both methods (5 mL for compendial vs. 400 µL for IMC), the CFU concentration is lower in the compendial method, potentially affecting its performance. On the other hand, as the maximum volume of the calScreener+ method is currently limited to 400 µL, this might in some cases complicate compliance with USP sample volume requirements. Possible approaches to meet the required volume could include dividing a sample over multiple vials, or using concentrated culture media, allowing larger sample volumes to be added.

Finally, while this study highlights the potential of IMC, further validation of the method as described in USP <1223> is required before implementation in GMP environments. This includes assessment of specificity, LOD, robustness, repeatability and ruggedness on a larger sample size (at least 75 to 100 samples), and on multiple inoculum levels.

## Conclusion

We assessed the performance of the calScreener+ IMC device using a diverse panel of microorganisms, consisting of reference strains and field isolates. Mean detection times were consistently lower for IMC than for the compendial USP <71> sterility test and IMC also detected a higher number of positive cultures. Our findings highlight IMC as a promising alternative to the compendial test, with the potential of speeding up batch releases while still detecting low-level contamination. Its continuous real-time readout and non-destructive nature makes it well-suited for implementation in GMP environments. Future research will be needed to validate the method, as well as assess calScreener+ performance in more complex matrices, such as cell-based products and formulations containing antimicrobial excipients.

## Supplementary materials

**Figure S1.**
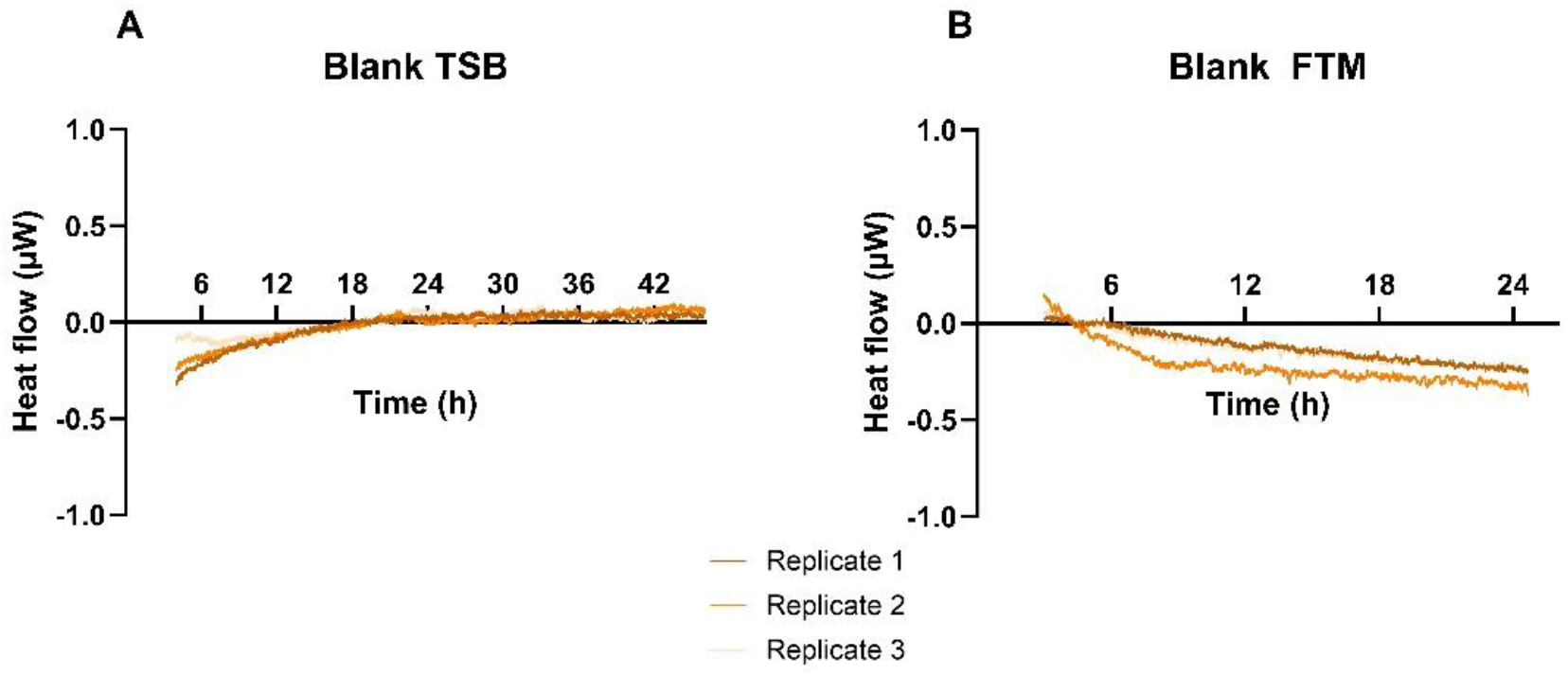
Thermograms of uninoculated samples of (A) TSB and (B) FTM, both in volumes of 400 µL. Short term noise and baseline drift equal 0.019 µW and 0.00069 µW/h for TSB and 0.041 µW and - 0.014 µW/h for FTM.

**Table S1.** Overview of all tested organisms with organism type and plate counts for 50 µL of suspension (mean ± standard deviation; n = 6).

| Organism | Organism type | Plate counts (CFU) |  |
| --- | --- | --- | --- |
|  |  | Aim 100 CFU | Aim 5 CFU |
| *Acidovorax delafieldii* LPM-S1 | Gram-negative | 30 ± 15 | 3 ± 2 |
| *Aspergillus brasiliensis* ATCC 16404 | Mold | 45 ± 22 | 3 ± 1 |
| *Aspergillus versicolor* LPM-S2 | Mold | 39 ± 10 | 3 ± 2 |
| *Bacillus spizizenii* ATCC 6633 | Gram-positive | 40 ± 12 | 2 ± 1 |
| *Bacillus subtilis* LPM-S3 | Gram-positive | 65 ± 18 | 2 ± 1 |
| *Burkholderia contaminans* LPM-S4 | Gram-negative | 65 ± 6 | 3 ± 2 |
| *Candida albicans* ATCC 10231 | Yeast | 52 ± 16 | 4 ± 2 |
| *Clostridium sporogenes* ATCC 19404 | Gram-positive | 37 ± 17 | 3 ± 2 |
| *Cutibacterium acnes* LMG 16711 | Gram-positive | 181 ± 49 | 11 ± 6 |
| *Micrococcus luteus* LPM-S5 | Gram-positive | 120 ± 24 | 7 ± 1 |
| *Penicillium chrysogenum* LPM-S6 | Mold | 55 ± 5 | 4 ± 2 |
| *Pseudomonas paraeruginosa* ATCC 9027 | Gram-negative | 72 ± 38 | 4 ± 2 |
| *Staphylococcus aureus* ATCC 6538 | Gram-positive | 89 ± 17 | 6 ± 3 |
| *Staphylococcus epidermidis* LPM-S7 | Gram-positive | 39 ± 15 | 3 ± 2 |
| *Staphylococcus hominis* LPM-S8 | Gram-positive | 36 ± 5 | 3 ± 2 |
| *Streptococcus pyogenes* LMG 14237 | Gram-positive | 22 ± 9 | 2 ± 1 |

